# Copulation calls indicate fertility but do not reflect female mate competition in wild Guinea baboons

**DOI:** 10.64898/2026.08.26.747217

**Authors:** Carolin Niederbremer, Federica Dal Pesco, Roger Mundry, Christof Neumann, Ndiouga Diakhate, Julia Fischer

## Abstract

Across different modalities, signals play a core role in attracting mates and influencing mating success. In several non-human primate species, females produce calls during mating that are thought to promote male competition over receptive females. The extent to which social system characteristics modulate the function of copulation calls remains less clear. We studied copulation calls in wild Guinea baboons (*Papio papio*), who live in a multilevel society structured around units in which females associate and mate almost exclusively with a single male. We hypothesised that females use copulation calls as an indirect form of mate competition, with competition increasing in larger units. In addition, we hypothesised that females are more likely to mate again after calling. We analysed 6116 copulations between 2014 and 2025, involving 99 reproductively active females and 78 subadult and adult males. Females produced copulation calls in 72.7% of copulations, with large inter-individual variation. Neither unit size nor its interaction with the female’s swelling size or the presence of simultaneously receptive females affected the probability of calling. A survival analysis with a subset of the data (2353 copulations) revealed no effect of calling on the latency to the next mating. Our results render the hypothesis that female Guinea baboons use calls in indirect mate competition unlikely. Yet, the probability of calling varied with sexual swelling size, suggesting that calls signal female fertility. Possibly, Guinea baboon copulation calls represent an evolutionary remnant, no longer under selective pressure, and can be considered index signals of female fertility.

## Introduction

Animals compete both intra- and intersexually for mating opportunities, often using elaborate traits called ornaments to attract the opposite sex (Rubenstein & Alcock, 2019). For example, in many bird species, males are equipped with colourful and/or elongated feathers which they present to females in courtship rituals, and females prefer mating with more conspicuous males (e.g., Goldie’s bird of paradise, *Paradisaea decora*; resplendent quetzal, *Pharomachrus mocinno*; Rubenstein & Alcock, 2019). Ornaments might be present in just one or both sexes, and if present in both sexes, ornamentation might be mutual (i.e., both sexes display the same ornaments) or differ between the sexes (Hare & Simmons, 2019). Ornaments might permanently advertise mate quality or emerge temporarily, for example in response to hormonal changes around the fertile window (Hare & Simmons, 2019), as illustrated by the reproductive colouration of female lagoon gobies (*Knipowitschia panizzae*) and alpine accentors (*Prunella collaris*; Massironi et al., 2005; Chiba & Nakamura, 2002). In the broadest sense, female signals that are linked to ovulation or motivation to mate can be conceived as ornaments.

In nonhuman primates, sexual swellings and copulation calls constitute the most conspicuous forms of female sexual signals (Clutton-Brock & Huchard, 2013). Copulation calls signal fertility in many species, as their acoustic structure may vary over the cycle and females are more likely to produce calls around the time of ovulation (for a review, see Pradhan et al., 2006). Copulation calls contain acoustic cues to caller identity (Semple, 2001) and may vary with other socially relevant information like whether ejaculation occurred (Belinchón et al., 2025; Engelhardt et al., 2012; Vaglio et al., 2020). Playback experiments showed that such variation is perceptually salient and affects male behaviour towards females (Pfefferle, Heistermann, et al., 2008).

A variety of functional explanations for copulation calls in female primates have been proposed, mostly focusing on promoting male competition over receptive females (reviewed in Pradhan et al., 2006). The male-male competition hypothesis postulates that females produce copulation calls to incite male competition, resulting in females mating with the most dominant males, as suggested for chacma baboons (*Papio ursinus*; Hamilton & Arrowood, 1978). The sperm competition hypothesis, in contrast, states that females produce copulation calls to attract and mate with as many males as possible, which best explains the use of copulation calls in Barbary macaques (Pfefferle, Brauch, et al., 2008). In contrast, the postcopulatory female choice hypothesis posits that females may produce copulation calls after mating with their preferred male to motivate mate guarding and thereby reduce sperm competition with unpreferred males (Maestripieri & Roney, 2005). Because competition for mates is also linked to social organisation, we expect sexual selection to operate differently in species living in different social systems, as for example in multi-level societies, in which females are associated with specific males, but still live in a group composed of multiple males and females. One such species is the Guinea baboon, which lives in a multi-level society with female-biased dispersal (Kopp et al., 2015; Patzelt et al., 2014). At the core of the multi-level society is the ‘unit’ comprising a primary male, one to eight females, and their young (Dal Pesco et al., 2022; Goffe et al., 2016; unpublished data). Females mate predominantly with their primary male, males rarely fight over females, and females may transfer relatively freely between males (Goffe et al., 2016). A previous study investigating copulation calls in captive Guinea baboons suggested that females call to incite mate-guarding behaviour and avoid the risk of sperm competition (Maestripieri et al., 2005).

To investigate the use of female copulation calls in Guinea baboons against the background of existing knowledge about copulation calls in other baboon species, we analysed copulations from 11 years of observational data recorded in a wild population in the Niokolo-Koba National Park in Senegal. We hypothesised that female Guinea baboons use copulation calls to attract male attention as a form of indirect mate competition among females within the same unit. This competition might be particularly pronounced if multiple females within a unit are simultaneously receptive and compete for mating opportunities. Further, a female’s motivation to compete might be higher during peak swelling, when she is most likely to conceive, compared to other swelling states.

We therefore predicted that females in larger units are more likely to produce copulation calls than those in smaller units. We also predicted that females are more likely to produce copulation calls when other females in the same unit are simultaneously receptive, and that they are more likely to do so at peak swelling than at other swelling states. In addition, females might depend on multiple copulations to increase the probability of conception, as some baboon species have a pattern of multiple intromissions to ejaculation (Dewsbury & Pierce Jr., 1989; Dixson, 2012), and giving a copulation call might increase the probability of being mated again (Semple, 1998). Therefore, we predicted that giving a copulation call increases the probability that a female will mate again with the same male. To address the previous observation that copulation calls incited mate guarding in a captive population (Maestripieri et al., 2005), we also tested whether males were more likely to follow a female depending on whether she called.

## Methods

### Study Subjects

The data collection for this study took place at the Centre de Recherche de Primatologie (CRP) Simenti in the Niokolo-Koba National Park in Senegal (Fischer et al., 2017) between April 2014 and June 2025. During this period, the Guinea baboon population (>400 individuals present simultaneously) around the field station comprised eight habituated and identified focal parties, although not all were present at the same time. Party size and composition varied during the study period (Table 1). Adult sex ratio calculations included all adult and subadult individuals for each party and year.

**Table 1.** Party size and composition of focal parties during the study period.

| Party | Observation period | Average number individuals | Range number individuals | Average adult sex ratio |
| --- | --- | --- | --- | --- |
| 9 | 2014 - 2020 | 44.5 | 37 to 51 | 0.96 |
| 6 | 2014 - 2020 | 39.9 | 35 to 48 | 1.01 |
| 5 | 2016 - 2025 | 36.9 | 28 to 45 | 0.70 |
| 9B | 2019 - 2025 | 18.4 | 6 to 28 | 0.70 |
| 6I | 2020 - 2025 | 17.5 | 11 to 25 | 1.12 |
| 6W | 2020 - 2025 | 25.1 | 20 to 32 | 0.97 |
| 13 | 2021 - 2025 | 29 | 15 to 34 | 0.58 |
| 15 | 2021 - 2023 | 6.2 | 3 to 9 | 0.53 |

### Data Collection

We recorded all data on handheld devices (Samsung Note 2 and Gigaset GX290) equipped with the Pendragon software (Pendragon Software Corporation, USA). We recorded copulations and noted whether the female produced a copulation call both during 20-minute focal-animal samples and ad libitum sampling (Altmann, 1974). We considered only copulations between subadult/adult females (N = 99 females) and subadult/adult males (N = 78 males; for age-category assessment and definitions, see Dal Pesco & Fischer, 2022) who belonged to our focal parties to ensure reliable information about unit composition and unit females’ reproductive states. We observed copulations between 350 different dyads. Only copulations with confirmed penetration were included in the analysis. If fresh sperm was visible on the hindquarters of the female or the penis of the male right after the copulation, we recorded the copulation as confirmed ejaculation. However, it is possible that in some of the copulations without confirmed ejaculation, males did ejaculate, but sperm was not clearly visible to the researchers.

We determined unit composition based on female-male interaction patterns (i.e., frequency of copulations, grooming bouts, contact-sit bouts, greetings, aggression events, duration of grooming and contact-sit bouts) (Dal Pesco et al., 2022), as it was previously shown that female Guinea baboons interact significantly more with their primary male (Goffe et al., 2016). We recorded the reproductive state of all females daily using morphological changes in the anogenital area and paracallosal skin. We categorised females as detumescent, tumescent, pregnant, or lactating (Dal Pesco & Fischer, 2022; Goffe et al., 2016; Table 2). For tumescent females, we distinguished between small, medium, and peak swellings (Dal Pesco & Fischer, 2022; Goffe et al., 2016; Table 2).

**Table 2.** Definitions of the reproductive states of female Guinea baboons at the CRP Simenti field site in the Niokolo-Koba National Park, Senegal (Dal Pesco & Fischer, 2022)

| Female reproductive state |  | Definition |
| --- | --- | --- |
| Detumescent |  | Absence of swelling of the anogenital area (AGA) or periccallosal skin (PCS). The skin around the anus and labia may be wrinkled and a bit loose post swelling or tight. |
| Tumescent | Small | A small swelling of the AGA. The swelling appears vertically and broadens slightly. |
|  | Medium | A medium-large swelling of the AGA and a small swelling of the PCS. The swollen area extends vertically and horizontally and comes outward slightly. |
|  | Peak | A large swelling of the AGA and a full swelling of the PCS. The swollen area is 3D often forcing the tail carriage to be higher and causing the female to sit slightly sideways; some areas protrude outward more than others. The width at peak swelling does not extend beyond the outer extremities of the ischial callosities in *Papio papio* as it does in other baboon species. |
| Pregnant |  | A tightness of the AGA and PCS and a change in skin colour to pink or red. The peak colour varies among females and the process begins with a slight pinkening of the AGA in most females. The AGA and PCS become bright pink over a number of weeks/months. In some females the labia may appear slightly swollen on and off throughout this period. |
| Lactating |  | Females with dependent offspring that have not started ovarian cycling again. The skin of the AGA will be tight but may be tinged red or pink on some females even a few months post parturition. |

### Statistical Analysis

We ran all analyses in R (version 4.5.1; R Core Team, 2025) and RStudio (version 2025.09.2; Posit team, 2025).

#### Effect of unit size on the probability of producing a copulation call

To estimate the extent to which the probability of producing a copulation call depended on the unit size (i.e., the number of subadult/adult females within a unit), we fitted a Generalized Linear Mixed Model (GLMM; Baayen, 2008) with binomial error structure and logit link function (McCullagh & Nelder, 1989) using the function “glmer” of the package lme4 (version 1.1-37; Bates et al., 2015). Our main predictor of interest was female unit size (count, range = 1-8). We included a female’s swelling size (factor with levels small, medium, peak) and whether at least one other co-resident female was tumescent (hereafter, co-resident female receptiveness) at the same time (factor with levels no, yes; if a female had no co-resident females, it was no by default). We included additional fixed effects to control for female age (factor with levels: subadult female, young adult female, mature adult female, old adult female) and male status (factor with levels: primary male, non-primary male) to indicate whether the female copulated with her primary male or with a different male. As we anticipated the effect of unit size to be greater in females with peak swellings, we included the interaction between unit size and swelling size. Furthermore, since we anticipated the effect of unit size to be more pronounced in the presence of other receptive females within the unit, we included the interaction between unit size and co-resident female receptiveness. We decided not to enter whether ejaculation occurred in our statistical analysis due to the potential bias from not reliably confirming ejaculation.

We included random intercepts for female identity (ID), male ID, as well as dyad ID, to account for their effects to avoid pseudoreplication, as we had multiple observations per individual and dyad. To keep the type 1 error rate at the nominal level of 0.05, we included all theoretically identifiable random slopes (Barr et al., 2013; Schielzeth & Forstmeier, 2009). We included random slopes for unit size, swelling size, co-resident female receptiveness, male status, the interaction between unit size and swelling size, and the interaction between unit size and co-resident female receptiveness, all within female ID. We included random slopes for unit size, swelling size, co-resident female receptiveness, female age, the interaction between unit size and swelling size, and the interaction between unit size and co-resident female receptiveness, all within male ID. We included correlations between random slopes and intercepts in the model (Barr et al., 2013).

Before fitting the model, we z-transformed unit size to a mean of zero and a standard deviation of one to ease model convergence and interpretation of model estimates (Schielzeth, 2010). We used parametric bootstrapping to determine fitted values and confidence intervals for model estimates (N=1000 bootstraps; function “bootMer” in the lme4 package; version 1.1-37; Bates et al., 2015). To assess model stability, we dropped the individual levels of the grouping factors one at a time (e.g., dropping each female ID once), fitted the full model to each subset, and compared the range of estimates with those from the full dataset. The estimates showed moderate to good stability (see Table 3 in the results section).

**Table 3.** Results of the full model of the effect of unit size on probability to produce copulation calls (estimates together with standard errors, confidence limits, significance tests, and the range of estimates after dropping levels of grouping factors one at a time)

| Term | Estimate | SE | Lower CI | Upper CI | $\chi^2$ | df | P | min | max |
| --- | --- | --- | --- | --- | --- | --- | --- | --- | --- |
| Intercept | 1.561 | 0.203 | 1.026 | 1.791 |  |  |  | 1.493 | 1.650 |
| Unit size<sup>(1)</sup> | 0.213 | 0.172 | -0.092 | 0.411 |  |  |  | 0.129 | 0.269 |
| Co-resident female receptiveness<sup>(2)</sup> | -0.058 | 0.166 | -0.302 | 0.163 |  |  |  | -0.154 | -0.002 |
| Small swelling<sup>(3)</sup> | -2.183 | 0.202 | -2.435 | -1.675 |  |  |  | -2.258 | -2.100 |
| Medium swelling<sup>(3)</sup> | -0.806 | 0.119 | -1.001 | -0.595 |  |  |  | -0.870 | -0.747 |
| Young adult female<sup>(4),(5)</sup> | -0.207 | 0.206 | -0.479 | 0.207 | 3.793 | 3 | 0.285 | -0.317 | -0.149 |
| Mature adult female<sup>(4),(5)</sup> | 0.015 | 0.21 | -0.199 | 0.606 |  |  |  | -0.096 | 0.113 |
| Old adult female<sup>(4),(5)</sup> | 0.585 | 0.373 | 0.318 | 1.668 |  |  |  | 0.383 | 0.917 |
| Male status<sup>(6)</sup> | -0.188 | 0.202 | -0.607 | 0.104 | 0.833 | 1 | 0.361 | -0.267 | -0.091 |
| Unit size<sup>(1)</sup>:Co-resident female receptiveness<sup>(2)</sup> | -0.169 | 0.172 | -0.334 | 0.174 | 0.965 | 1 | 0.326 | -0.239 | -0.074 |
| Unit size<sup>(1)</sup>:Small<br>swelling<sup>(3),(7)</sup> | -0.072 | 0.201 | -0.307 | 0.359 | 1.113 | 2 | 0.573 | -0.162 | 0.011 |
| Unit<br>size<sup>(1)</sup>:Medium<br>swelling<sup>(3),(7)</sup> | 0.108 | 0.129 | -0.156 | 0.329 |  |  |  | 0.034 | 0.166 |

<sup>(1)</sup> z-transformed to a mean of zero and a standard deviation (sd) of one; mean and sd of the original variable were 3.14 and 1.72, respectively
<sup>(2)</sup> dummy coded with no being the reference level
<sup>(3)</sup> dummy coded with peak swelling being the reference level
<sup>(4)</sup> dummy coded with subadult female being the reference level
<sup>(5)</sup> the indicated test refers to the overall effect of female age category
<sup>(6)</sup> dummy coded with primary male being the reference level
<sup>(7)</sup> the indicated test refers to the overall effect of the interaction between unit size and swelling size

As an overall test of the fixed effects of unit size and its interactions with swelling size and co-resident female receptiveness, and to avoid cryptic multiple testing (Forstmeier & Schielzeth, 2011), we conducted a full-null model comparison. The null model lacked unit size and, hence, also the interactions in the fixed-effects part, but was otherwise identical. We used likelihood ratio tests to assess the significance of individual fixed effects (Dobson & Barnett, 2018) by comparing the full model with models lacking one fixed effect at a time (R function “drop1”). The sample analysed with this model included 6116 copulations among 99 females and 78 males, comprising 350 dyads. Copulation calls occurred during 4448 of the copulations.

#### Effect of copulation calls on the latency to copulate again

With a second analysis, we investigated whether two individuals are more likely to copulate again within a given focal observation period if the female produced a copulation call compared to instances when no copulation call occurred. We fitted a Bayesian regression model with a Weibull error distribution and log link function using the “brm” function in the brms package (version 2.23.0; Bürkner, 2018). We used the default priors of brms.

To test whether the occurrence of a copulation call influenced the latency to the following copulation, we only considered copulations recorded during focal observations. We calculated the time in seconds from the onset of the first observed copulation until either another copulation between the same male and female occurred or the focal observation was interrupted (due to out of sight) or ended. This analysis included 2353 copulations. We excluded two copulations that occurred exactly when the respective focal observation ended, resulting in a time until the end of the observation period of zero. We noted censor status (factor with levels: no, yes) for each copulation and included it in the response. If more than two copulations took place within a focal observation, we calculated latencies between the current copulation and the preceding one for all respective copulations. The main predictor of interest was whether a copulation call occurred during the preceding copulation (levels: no, yes). As control predictors, we included all fixed effects from the binomial model but without interactions, with their respective levels: unit size, co-resident female receptiveness, swelling size, female age, and male status.

We included random intercept effects for female ID, male ID, and dyad ID and all theoretically identifiable random slopes. These included random slopes for copulation call, unit size, co-resident female receptiveness, and swelling size within female ID and copulation call, unit size, swelling size, and female age within male ID. We included correlations between random slopes and intercepts in the model. Before fitting the model, we z-transformed unit size to a mean of zero and a standard deviation of one to ease model convergence and interpretation of model estimates (Schielzeth, 2010). The model was based on 2353 copulations from 1424 focal samples, involving 94 females and 74 males, comprising 268 dyads. 816 copulations were followed by another copulation, and the observation periods of 1537 copulations were censored.

#### Effect of copulation calls and swelling size on male-initiated proximity

To test whether female copulation calls increase the probability of reestablishing proximity after separation, we considered only events recorded during focal observations in which the female left the male immediately after the copulation. We recorded whether the male approached the female at least once during the observation period after the separation (binary response: no, yes). We included the interaction between copulation call (levels: no, yes) and swelling size (levels: small, medium, peak) in our model. The observation period lasted up to 10 minutes, following Maestripieri et al. (2005), but ended sooner if the focal observation ended, the animal went out of sight, or another copulation occurred between the same male and female (average observation period: 5.5 min). As the probability of observing any behaviour increases with observation effort, we fitted a non-linear model to account for varying observation duration:

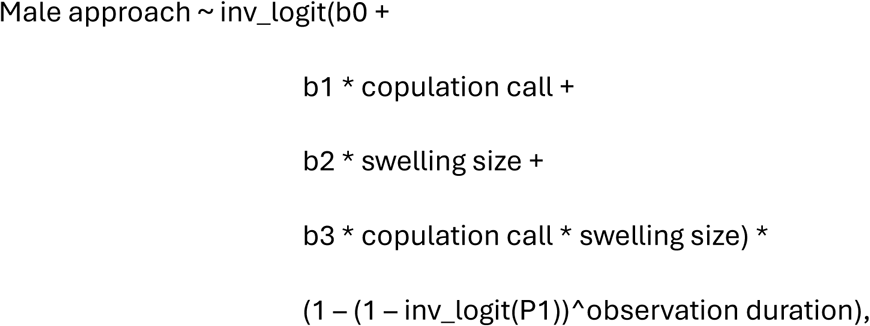

where b0 is the intercept term, b1 is the slope estimate for copulation call, b2 is the slope estimate for swelling size, b3 is the slope estimate for the interaction term, and P1 is the estimate for the probability to observe an approach in one unit of observation time.

We fitted the model using the “brm” function with Bernoulli error distribution with identity link function in the brms package (version 2.23.0; Bürkner, 2018). We used Normal(0, 1) priors for the intercept and main effects, while we set a tighter Normal(0, 0.5) prior for the interaction term. For p1 we set a Normal(−1, 1) prior. For variance estimates we used brms’ default priors.

We included random intercept effects for female ID, male ID, dyad ID, and the combination of dyad and session ID, as well as all theoretically identifiable random slopes. These included random slopes for copulation call and swelling size within female ID, and copulation call, swelling size, and their interaction within male ID. The model was based on 1952 copulations from 1217 focal samples, involving 92 females and 73 males, comprising 257 dyads. Males approached the female again after 511 copulations.

### Ethical Note

We followed all applicable international, national, and/or institutional guidelines for the care and use of animals. Research was conducted in accordance with the regulations set by the Senegalese agencies and the ethical standards of the Animal Care Committee at the German Primate Centre. This study adhered to the ASAB guidelines for the ethical treatment of nonhuman animals in behavioural research (ASAB Ethical Committee/ABS Animal Care Committee 2023). Approval and research permission were granted by the DPN and the MEPN de la République du Sénégal.

## Results

Females in our sample produced copulation calls in 4448 out of 6116 (72.7%) of copulations. However, there was large individual variation in the probability of producing copulation calls (see Table S1 for estimated random-effects parameters). Considering the whole dataset (N = 99 females; range of copulations per female: 1–247), the proportion of copulations with copulation calls varied between 0% and 100% for individual females. This variation did not change when we excluded potential small-sample effects by considering only females who copulated at least 40 times during our study period (N = 58 females). In this sample, the proportion of copulations with a copulation call ranged from 10.8% to 96.7% for individual females. The proportion of copulations with calling was similar during focal animal samples (N = 2355 copulations, 70.7% with copulation call) and ad libitum sampling (N = 3761 copulations, 74.0% with copulation call). The proportion of copulation calls given during copulation with confirmed ejaculation (N = 645 copulations, 79.2% with copulation call) was similar to the proportion of copulation calls given during copulations when we could not confirm ejaculation (N = 5471 copulations, 72.0% with copulation call).

The full-null model comparison showed that neither unit size, nor the interaction between unit size and co-resident female receptiveness (Figure 1) or that between unit size and swelling size of the copulating female (Figure S1) had a statistically significant effect on call probability (full-null model comparison: χ^2^_4_ = 4.34, P = 0.362; Table 3). Thus, we fitted a reduced model that excluded the interactions between unit size and swelling size, as well as the interaction between unit size and co-resident female receptiveness, but was otherwise identical (Table S2). The reduced model also showed that unit size alone did not significantly affect call probability (Estimate ± SE = 0.19 ± 0.12, P = 0.105; Figure 2; Table S2). However, we found that the size of the copulating female’s swelling was significantly associated with call probability, with females with peak swellings being more likely to call (likelihood ratio tests of the overall effect of swelling size: χ^2^_2_= 64.99, P < 0.001; Figure 3; Table S2). Females at peak swelling were about twice as likely to produce copulation calls as females with small swellings (probability to call at small swelling: 34%, at medium swelling: 68%, at peak swelling: 83%; Figure 3; Table S2). Male status did not affect calling probability: females who copulated with their primary male were nearly as likely to give a copulation call (73.3% of the cases) as females who copulated with a male who was not their primary male (68% of the cases; P = 0.353; Table 3).

**Figure 1.**
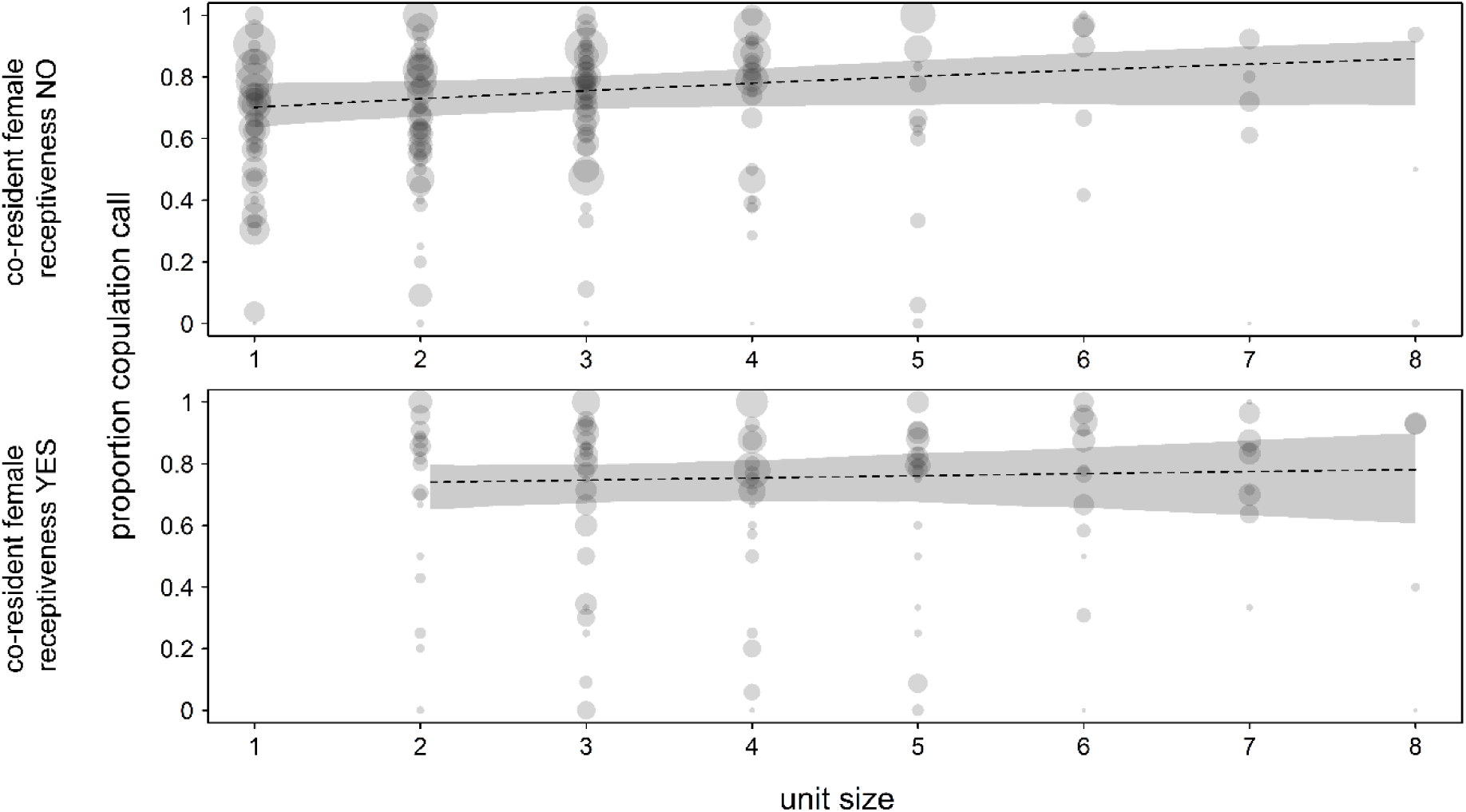
Proportion of copulation calls given in relation to unit size and co-resident female receptiveness. Indicated are the fitted model (dashed lines) and its 95% confidence intervals (grey polygons) with all other terms in the model being at their average. Dots represent the proportion of copulations with calls for each combination of unit size and female ID. The area of the dots is proportional to the number of copulations per female and unit size (range: 1 to 110).

**Figure 2.**
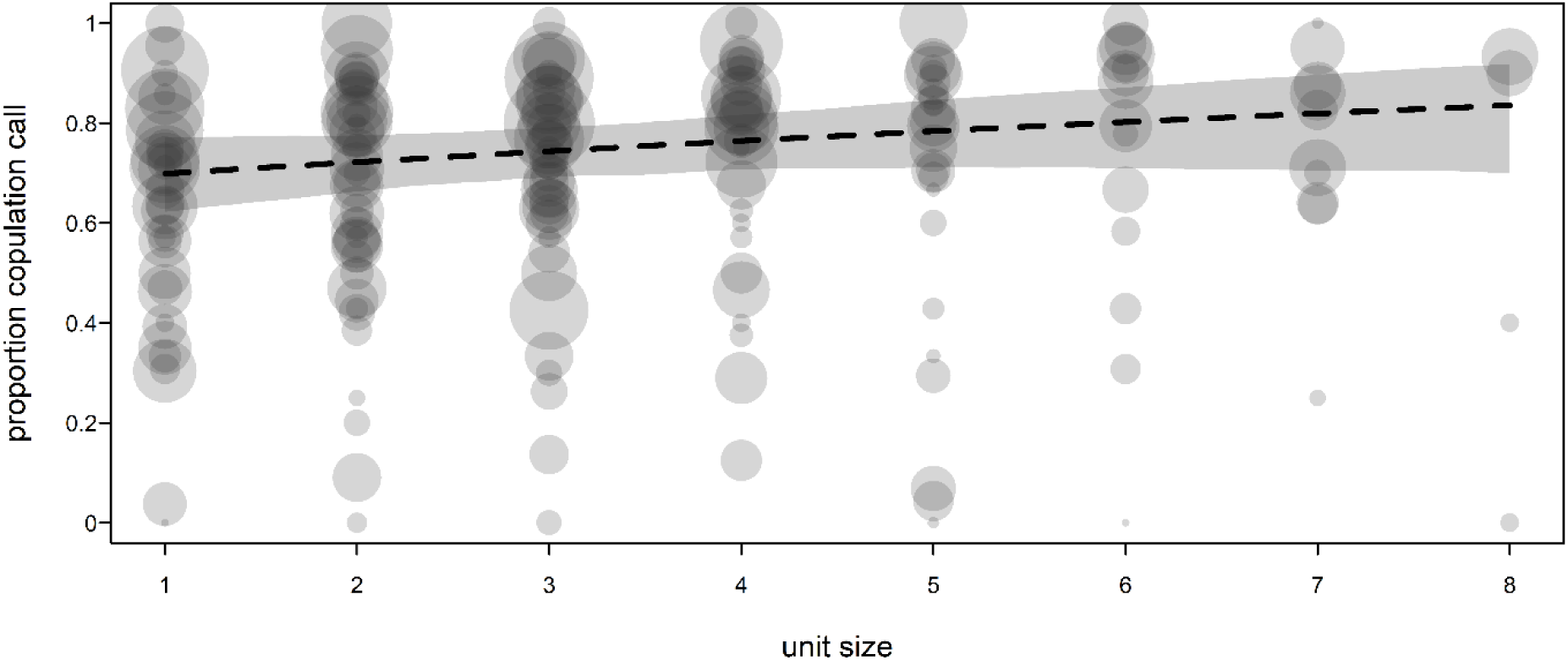
Proportion of copulation calls given in relation to unit size (reduced model). Indicated are the fitted model (dashed line) and its 95% confidence intervals (grey polygons) with all other terms in the model being at their average. Dots represent the proportion of copulations with calls for each combination of unit size and female ID. The area of the dots is proportional to the number of copulations per female and unit size (range: 1 to 110).

**Figure 3.**
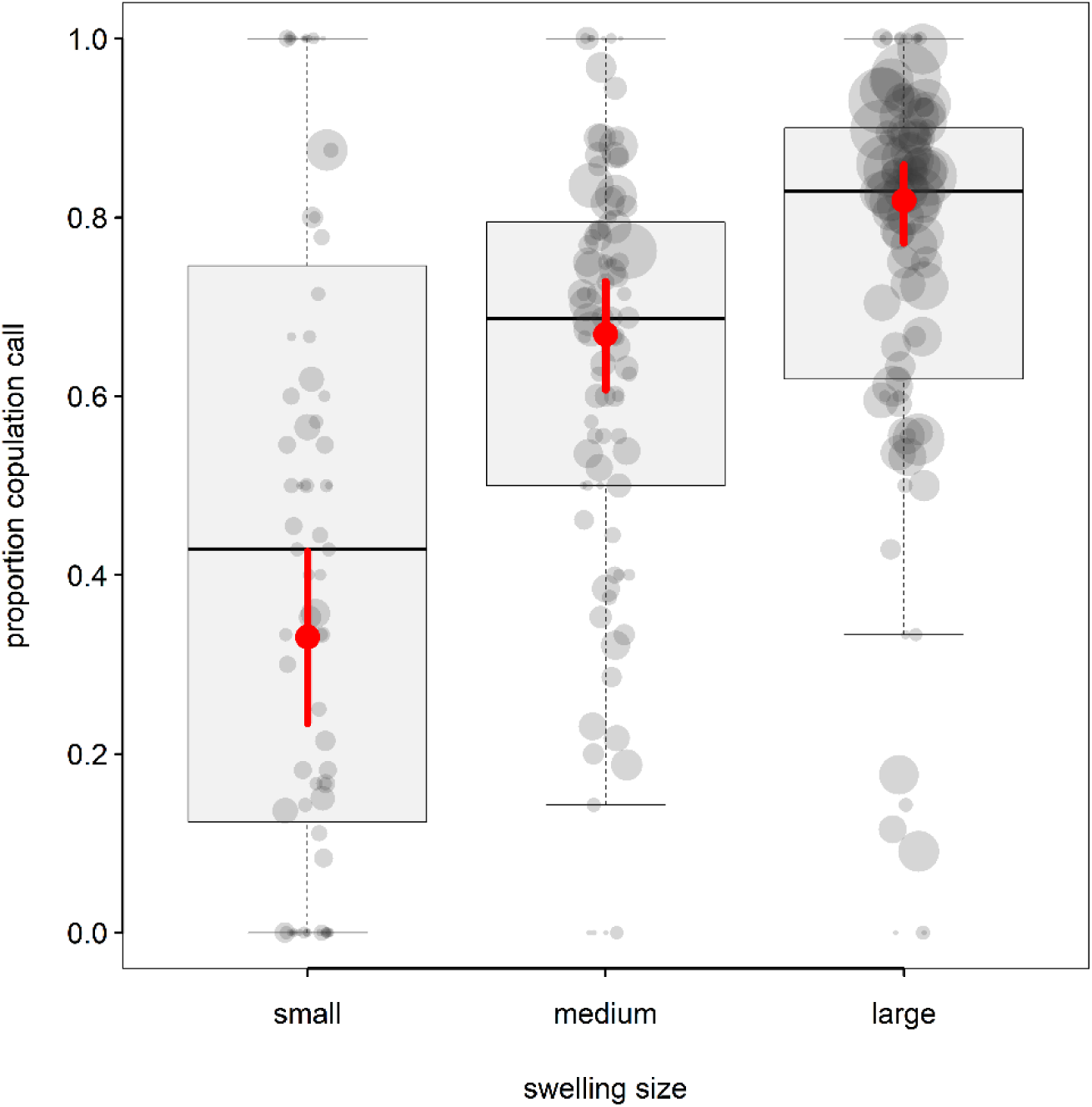
Proportion of copulation calls given in relation to swelling size. Indicated for each swelling size are the model predictions (red dots) with confidence intervals (red vertical lines) with all other terms in the model being at their average, median (horizontal line within each box), interquartile range (box), and 1.5 x the interquartile range (whiskers). Dots represent the proportion of copulations with calls for each combination of swelling size and female ID. Dot size (area) is proportional to the number of copulations per female and swelling size (range: 1 to 161).

Females produced copulation calls in 70.7% of the copulations included in the survival model (N = 2353 copulations). Most observation periods after copulations (65.3%) were censored, and females did not copulate again with the same male within the observation period. Producing a copulation call did not affect the probability of copulating again (Table 4). When females did not produce a copulation call, the initial copulation was followed by another one in 30.7% of the time, with a mean latency of 255 ± 205 s (range: 2-1051 s, N = 211). In cases when females did produce a copulation call, the copulation was followed by another copulation in 36.3% of the time, with a mean latency of 312 ± 214 s (range: 2-1110 s, N = 605). These differences were not statistically significant (Table 4). Overall, males approached the female again after 26.2% of copulations. We found no clear evidence that calling affected the probability that a male re-established contact with the female (Table 5; Figure 4).

**Figure 4.**
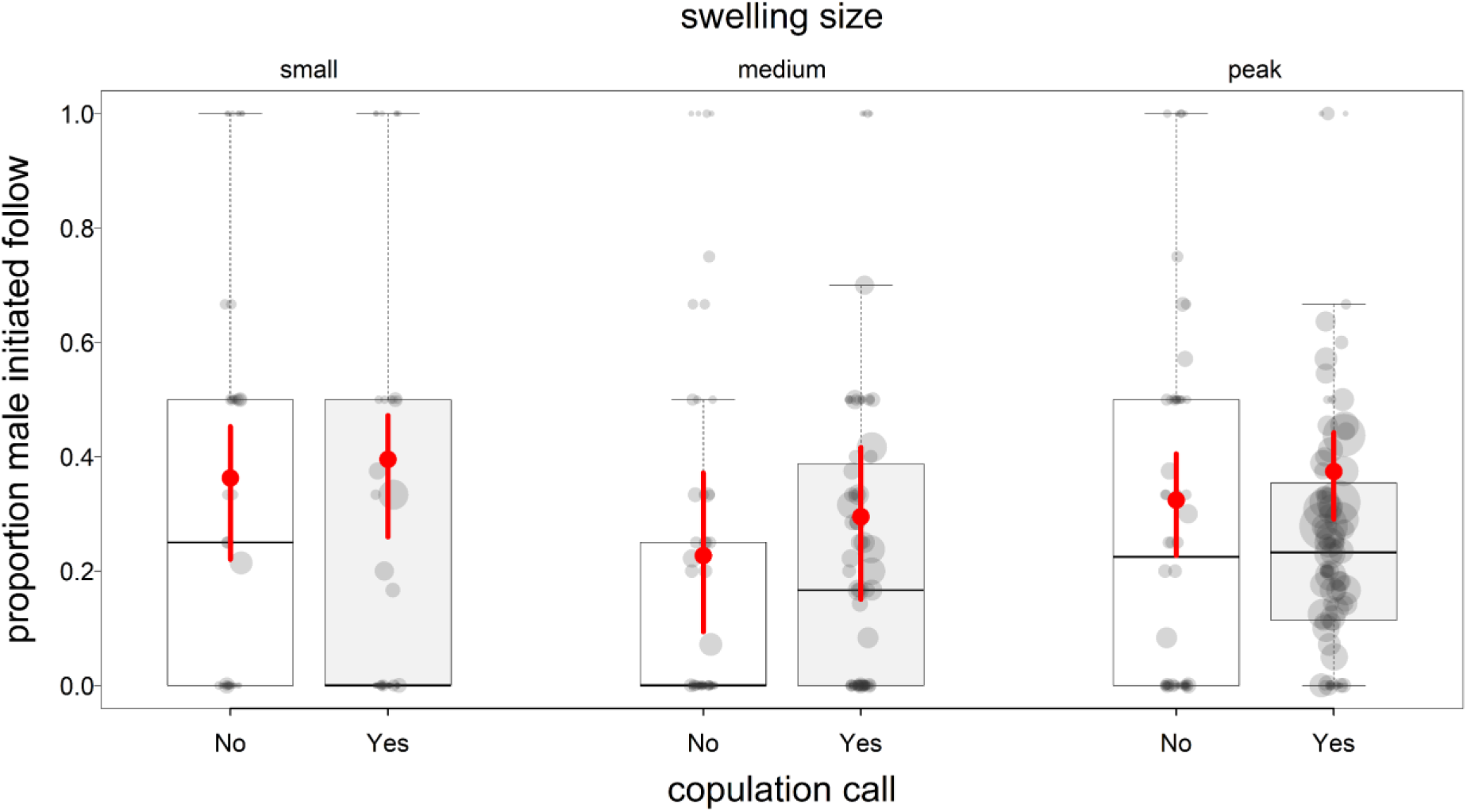
Proportion of copulations after which the male approached the female in relation to the combination of swelling size and copulation call. Indicated for each combination of swelling size and copulation call are the model predictions (red dots) with credible intervals (red vertical lines), median (horizontal line within each box), interquartile range (box), and 1.5 x the interquartile range (whiskers). Grey dots represent the observed proportion of copulations with male follow for each combination of swelling size, copulation call, and female ID. The area of the dots is proportional to the number of copulations per female and combination of copulation call and swelling size (range: 1 to 61).

**Table 4.**
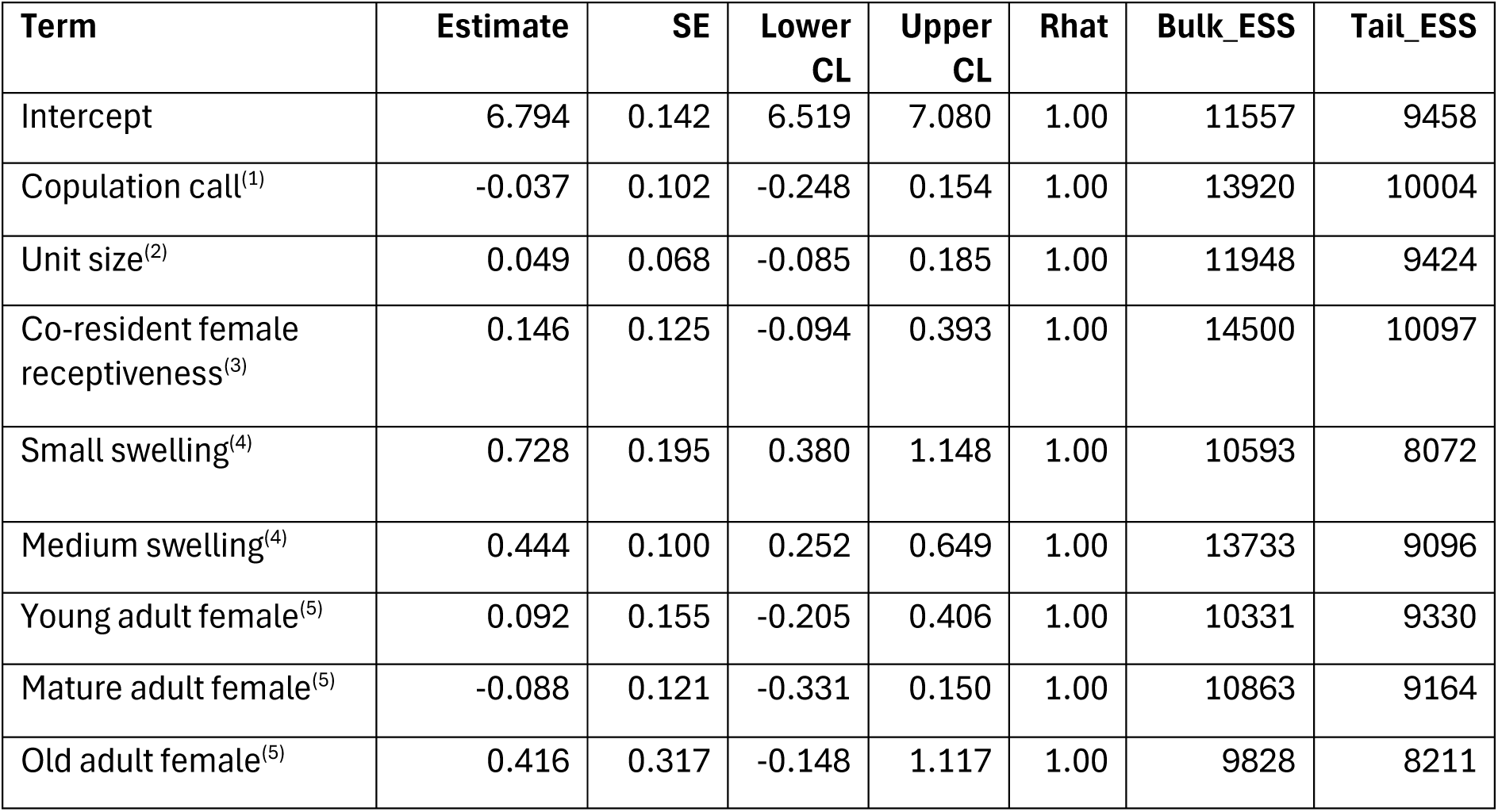
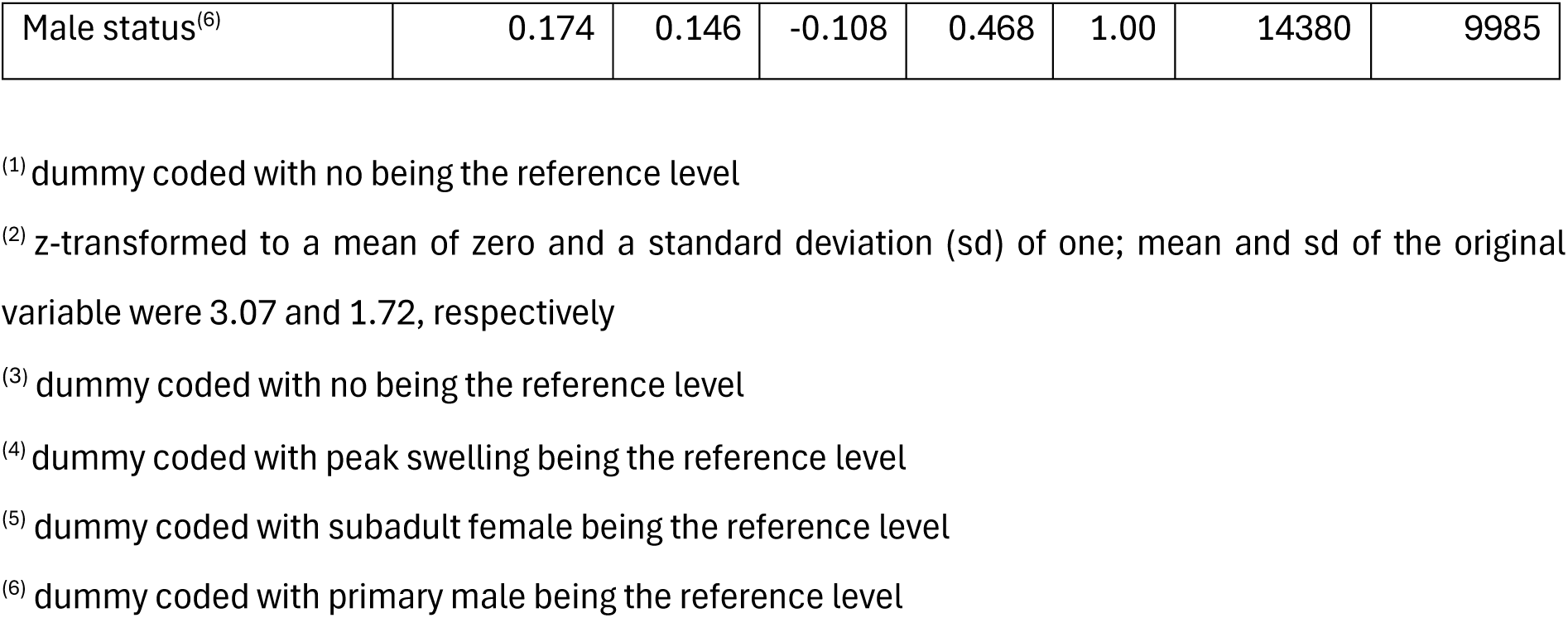
Results of the survival model of the effect of copulation call on the probability to copulate again (estimates together with standard errors, 95% credible limits, Rhat, and the effective sample size measures for each parameter)

**Table 5.** Results of the male-initiated proximity model of the effect of copulation call and swelling size on the probability of male approaches (estimates together with standard errors, 95% credible limits, Rhat, and the effective sample size measures for each parameter)

| Term | Estimate | SE | Lower CL | Upper CL | Rhat | Bulk_ESS | Tail_ESS |
| --- | --- | --- | --- | --- | --- | --- | --- |
| Intercept | 0.886 | 0.657 | -0.293 | 2.232 | 1.00 | 1544 | 2298 |
| Copulation call<sup>(1)</sup> | 0.671 | 0.632 | -0.496 | 1.952 | 1.00 | 2156 | 2692 |
| Small swelling<sup>(2)</sup> | 0.579 | 0.817 | -1.030 | 2.146 | 1.00 | 3503 | 2890 |
| Medium swelling<sup>(2)</sup> | -0.926 | 0.700 | -2.291 | 0.502 | 1.00 | 3211 | 2632 |
| Copulation call<sup>(1)</sup>:Small swelling<sup>(2)</sup> | -0.010 | 0.495 | -0.963 | 0.995 | 1.00 | 6472 | 3011 |
| Copulation call<sup>(1)</sup>:Medium swelling<sup>(2)</sup> | 0.052 | 0.457 | -0.826 | 0.932 | 1.00 | 4326 | 2846 |
| P1<sup>(3)</sup> | -2.095 | 0.119 | -2.323 | -1.846 | 1.00 | 591 | 1005 |

<sup>(1)</sup> dummy coded with no being the reference level
<sup>(2)</sup> dummy coded with peak swelling being the reference level
<sup>(3)</sup> estimated probability of a male approach within one unit observation time

## Discussion

We investigated the extent to which wild female Guinea baboon copulation calls play a role in mate competition, using a large dataset including more than 6000 copulations involving 350 different dyads. Females produced copulation calls in most copulations, but calling varied substantially among individuals. Contrary to our prediction, neither unit size nor its interactions with female swelling size or the presence of receptive co-resident females affected a female’s probability of giving a copulation call to a substantial degree, providing no support for the hypothesis that females use copulation calls to compete for their primary male’s attention. Large units with more than five females are rare in our study population (of 60 primary males, four had at some point during the study period six or more females), and we think that the differences detected with the reduced model are too small and uncertain to be of practical relevance to females. We also found no evidence that calling increased the probability of mating again with the same male, as calling did not affect the latency to the next mating, nor did calling affect males’ probability of reestablishing contact with females after they left after copulation. We therefore deem the postcopulatory female choice hypothesis unlikely. Instead, calling probability increased with female swelling size, suggesting that copulation calls may be linked to signalling female reproductive state rather than inciting mate competition.

Copulation calls have been reported in most baboon species (Table 5) and can likely be considered an ancestral trait. Many other Papionins (e.g., *Macaca*: long-tailed macaques: Engelhardt et al., 2012; Barbary macaques: Pfefferle, Brauch, et al., 2008; *Theropithecus*: gelada, *T. gelada*: Zanoli et al., 2022; *Lophocebus*: grey-cheeked mangabey, *L. albigena johnstoni*: Arlet et al., 2007; *Cercocebus*: Sooty mangabey, *C. torquatus atys*: Range & Fischer, 2004) likewise produce copulation calls. However, the frequency of calling and the prevalence of individual differences vary substantially among species, potentially influenced by a species’ mating system and the resulting strength of selective pressure on this trait.

Differences in the mating system are reflected in the evolutionary history of baboons: the genus *Papio* originated in southern Africa and diverged into a southern lineage (today chacma baboon, yellow baboon, *P. cynocephalus*, and Kinda baboon, *P. kindae*) and a northern lineage (today olive baboon, *P. anubis,* hamadryas baboon, *P. hamadryas*, and Guinea baboon; Fischer et al., 2019). Similar to the hypothesised ancestral form, chacma, olive, Kinda, and yellow baboons (‘COKY baboon’; Jolly, 2020) live in multi-male multi-female groups, and females can mate with several males during fertile periods. Hamadryas and Guinea baboons, in contrast, live in multi-level societies in which females are associated with a single male and mate predominantly with him (Fischer et al., 2019). The degree of male competition follows a gradient from south to northwest (Barrett & Henzi, 2008) and is very pronounced in the southern chacma baboon, whereas Guinea baboons show high degrees of tolerance and comparably little overt aggression (Dal Pesco & Fischer, 2020; Kalbitzer et al., 2015, 2016).

Copulation calls have been hypothesized to promote male competition over receptive females (Pradhan et al., 2006), and evidence from both captive and wild studies in the genus *Papio* supports a link between the degree of male competition and the prevalence of copulation calls (Table 6), with very high calling frequency and little individual variation in chacma baboons compared to lower calling frequency and very high individual variation in hamadryas and Guinea baboons (Table 6). Large individual variation suggests only weak selective pressure on call production, which in turn fits with the mating systems of both Guinea and hamadryas baboons. Vocalisations whose likely ancestral function was to incite male-male competition over receptive females (Pradhan et al., 2006) seem to have lost this purpose in the context of sexual selection in these species.

**Table 6.**
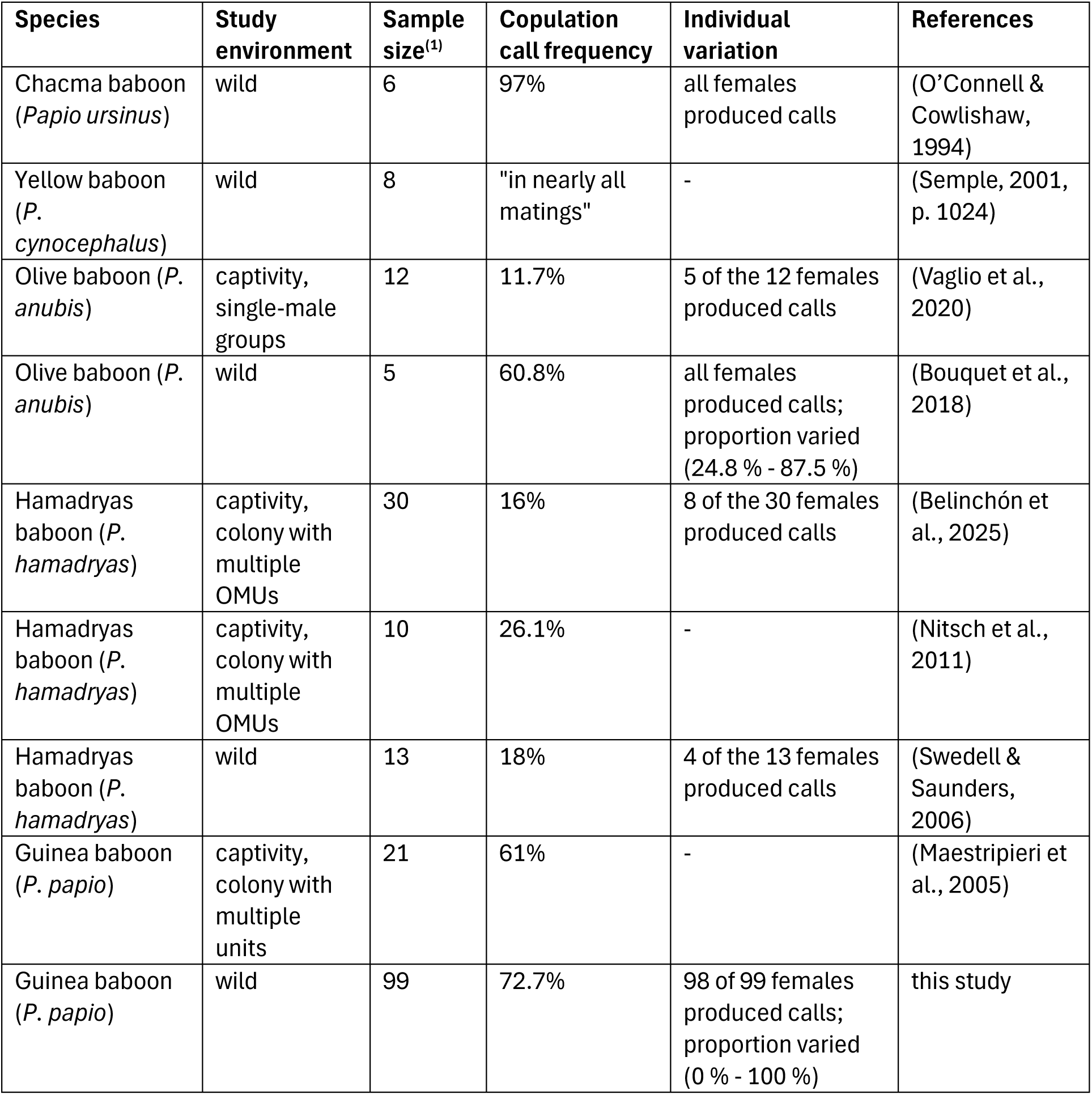
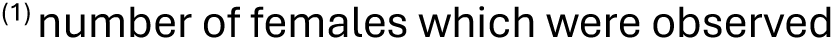
Comparison of female copulation call production in species of the genus *Papio*.

| Species | Study environment | Sample size<sup>(1)</sup> | Copulation call frequency | Individual variation | References |
| --- | --- | --- | --- | --- | --- |
| Chacma baboon (*Papio ursinus*) | wild | 6 | 97% | all females produced calls | (O'Connell & Cowlshaw, 1994) |
| Yellow baboon (*P. cynocephalus*) | wild | 8 | "in nearly all matings" | - | (Semple, 2001, p. 1024) |
| Olive baboon (*P. anubis*) | captivity, single-male groups | 12 | 11.7% | 5 of the 12 females produced calls | (Vaglio et al., 2020) |
| Olive baboon (*P. anubis*) | wild | 5 | 60.8% | all females produced calls; proportion varied (24.8 % - 87.5 %) | (Bouquet et al., 2018) |
| Hamadryas baboon (*P. hamadryas*) | captivity, colony with multiple OMUs | 30 | 16% | 8 of the 30 females produced calls | (Belinchón et al., 2025) |
| Hamadryas baboon (*P. hamadryas*) | captivity, colony with multiple OMUs | 10 | 26.1% | - | (Nitsch et al., 2011) |
| Hamadryas baboon (*P. hamadryas*) | wild | 13 | 18% | 4 of the 13 females produced calls | (Swedell & Saunders, 2006) |
| Guinea baboon (*P. papio*) | captivity, colony with multiple units | 21 | 61% | - | (Maestripieri et al., 2005) |
| Guinea baboon (*P. papio*) | wild | 99 | 72.7% | 98 of 99 females produced calls; proportion varied (0 % - 100 %) | this study |

<sup>(1)</sup> number of females which were observed

Without selective pressures specifically favouring copulation call production, females might indeed benefit from suppressing calls to prevent potential postcopulatory aggression by concealing certain copulations (Bouquet et al., 2018). Aggression by third parties in the mating context has been reported in several primate species (reviewed in Niemeyer & Anderson, 1983; e.g., eastern chimpanzee, *Pan troglodytes schweinfurthii*: Townsend et al., 2008), and female Guinea baboons might be subjected to male aggression if they copulate with males other than their primary male. Accordingly, females should be less likely to call when copulating with males other than their primary male to avoid risking aggression from their primary male. However, in our sample, the probability of females producing a copulation call was not related to male status, and females were as likely to call after copulating with their primary male as with any other male. We therefore do not conclude that the relatively low-amplitude calling in Guinea baboon females results from avoiding aggression from third parties.

If copulation calls in Guinea baboons neither present an obvious advantage nor disadvantage, the question remains why females produce them. However, an adaptation that was beneficial in the past might persist even after losing its function (Shanahan, 2011). Several vestigial behaviours and characteristics, traits that were initially maintained by selection but have become non-adaptive, are documented in the animal kingdom (Fong et al., 1995; Rayner et al., 2022). Male courtship and mating behaviours in several arthropods, for example, were retained after transition to asexuality and loss of their function (van der Kooi & Schwander, 2014). While reversed selection might result in trait loss, traits that are not costly to maintain and thus under relaxed selection might persist even without fitness benefits (Rayner et al., 2022). However, the expression of non-adaptive traits might be reduced over time (Fong et al., 1995). A reduction of trait expression, namely reduced salience in copulation calls of hamadryas and Guinea baboons, becomes visible when comparing their copulation call spectrograms with those of COKY baboons (Swedell & Saunders, 2006, p. 32). Swedell and Saunders (2006) report that copulation calls of hamadryas baboon females are quieter, shorter, and less complex in comparison to copulation calls of chacma and yellow baboons.

Spectrograms of Guinea baboon copulation calls (Kemp et al., 2017, p. 42; Maciej et al., 2013) resemble those of hamadryas baboons, and calls are shorter and quieter compared to chacma baboons (Dixson, 2012). Despite differences in amplitude and call length as observed for copulation calls, the structure of vocalisations in baboons seems to be, in general, highly conserved (Hammerschmidt & Fischer, 2019), and we assume that the underlying pattern generator is the same across baboon species. A similar picture arises when comparing ritualised greetings among baboon species (Dal Pesco & Fischer, 2020). While greetings are present in all baboon species, their frequency, form, and function vary with the degree of mate competition between species (Dal Pesco & Fischer, 2020). Signals thus might change in amplitude and function between species rather than disappearing or being replaced by completely new signals.

In our sample, female swelling size strongly predicted the probability of producing a copulation call, with females being twice as likely to produce a copulation call at peak swelling as at small swelling. Swelling size can be used as a proxy for fertility, as it varies across the cycle and reaches its maximum around ovulation (Brauch et al., 2007; Higham et al., 2008; Pfefferle et al., 2011; Street et al., 2016). The correlation between calling probability and swelling size indicates that both are under hormonal control and raises the possibility that females call to advertise their fertility. Consistent with this interpretation, studies in other primates likewise report a higher probability of calling during peak swelling than during other swelling states (Belinchón et al., 2025; Clay et al., 2011; Rigaill et al., 2013). Variation in calling may also be mediated by variation in sensitivity of the anogenital area, which, in turn, is under hormonal control (Henzi, 1996; Semple et al., 2002).

In summary, female Guinea baboon copulation calls seem to be closely linked to the female’s reproductive state, while they have largely lost their functional significance in the context of sexual selection. The prevalence of copulation calls without fulfilling an obvious function in the mating context, namely neither inciting male-male competition nor reflecting female competition over mates, suggests that their current persistence may not have a direct adaptive benefit. This finding illustrates that caution is needed when viewing behaviour through an adaptationists’ lens. While it is often assumed that behaviours must be adaptive to persist, our study shows that, in the absence of obvious costs associated with reversed selection, traits that do not benefit their bearer might persist or only slowly degrade (Rayner et al., 2022).

## Acknowledgements

We are grateful to the Direction des Parcs Nationaux (DPN) and the Ministère de l’Environnement et de la Protection de la Nature (MEPN) de la République du Sénégal for allowing us to work in the Parc National du Niokolo-Koba (PNNK). We particularly thank all the Conservateurs of the park during the study period (Oussoumane Kane, Mallé Gueye, Amar Fall, Assane Ndoye, Jacques Gomis, Paul Moïse Diedhiou, and Ibrahima Ndao), as well as the deputy Conservateur, Elhadji Mamadou Thiaw, for their cooperation and support. We thank all the CRP Simenti field assistants and students, in particular Cheikh Sané, Moustapha Dieng, Moustapha Faye, Armel Louis Nyafouna, El’Hadji Yankhoba Dansokho, Touradou Sonko, Vieux Biaye, Djibril Coly, Amadou Bamba Diedhiou, Chérif Younousse Kéba Camara, Jean Louis Diouf, Lassana Ba, Jean Malack, Boubacar Sow, Malamine Diedhiou, Moussa Dieng, Dame Diallo, Hamidou Sogoba, and Alfred Oga, for their support and work in the field. This research was funded by the Deutsche Forschungsgemeinschaft (DFG, German Research Foundation) under Project-ID 454648639 SFB 1528 “Cognition of Interaction” and Project-ID 254142454/GRK 2070 “Understanding Social Relationships” and by the Leibniz ScienceCampus “Primate Cognition”.

## Author Contributions

Conceptualization: CN, FDP, JF; Methodology: CN, FDP, RM, ChN, JF; Formal analysis: CN, FDP, RM, ChN; Investigation: CN, FDP, ND; Data curation: FDP; Writing – Original Draft: CN; Writing – Review & Editing: CN, FDP, RM, ChN, ND, JF; Visualization: CN, FDP, RM, ChN, JF; Funding acquisition: JF.

## Data Availability Statement

The data and code for the statistical analysis that support the findings will be made publicly available on GRO upon acceptance of the manuscript.

## Notes

### Competing Interest Statement

The authors have declared no competing interest.

