## supplemental Table 1 and 2 and supplemental Figure 1 for "Copulation calls indicate fertility but do not reflect female mate competition in wild Guinea baboons"

**Supplementary Material**

**Table S1.** Estimated random effects parameters of the full model of the effect of unit size on the probability to produce copulation calls.

| **Grouping factor** | **Term 1** | **Term 2** | **SD / correlation** |
| --- | --- | --- | --- |
| Dyad ID | Intercept | - | 0.644 |
| Female ID | Intercept | - | 0.808 |
| Female ID | Unit size^(1)^ | - | 0.464 |
| Female ID | Co-resident female receptiveness^(2)^ | - | 0.296 |
| Female ID | Small swelling^(3)^ | - | 0.711 |
| Female ID | Medium swelling^(3)^ | - | 0.440 |
| Female ID | Male status^(4)^ | - | 0.725 |
| Female ID | Unit size^(1)^:Co-resident female receptiveness^(2)^ | - | 0.383 |
| Female ID | Unit size^(1)^:Small swelling^(3)^ | - | 0.549 |
| Female ID | Unit size^(1)^:Medium swelling^(3)^ | - | 0.425 |
| Female ID | Intercept | Unit size^(1)^ | 0.375 |
| Female ID | Intercept | Co-resident female receptiveness^(2)^ | 0.948 |
| Female ID | Intercept | Small swelling^(3)^ | -0.034 |
| Female ID | Intercept | Medium swelling^(3)^ | -0.670 |
| Female ID | Intercept | Male status^(4)^ | -0.428 |
| Female ID | Intercept | Unit size^(1)^:Co-resident female receptiveness ^(2)^ | -0.859 |
| Female ID | Intercept | Unit size^(1)^:Small swelling ^(3)^ | -0.652 |
| Female ID | Intercept | Unit size^(1)^:Medium swelling ^(3)^ | -0.484 |
| Female ID | Unit size^(1)^ | Co-resident female receptiveness^(2)^ | 0.472 |
| Female ID | Unit size^(1)^ | Small swelling^(3)^ | 0.129 |
| Female ID | Unit size^(1)^ | Medium swelling^(3)^ | -0.253 |
| Female ID | Unit size^(1)^ | Male status^(4)^ | 0.125 |
| Female ID | Unit size^(1)^ | Unit size^(1)^:Co-resident female receptiveness^(2)^ | -0.740 |
| Female ID | Unit size^(1)^ | Unit size^(1)^:Small swelling^(3)^ | -0.195 |
| Female ID | Unit size^(1)^ | Unit size^(1)^:Medium swelling^(3)^ | -0.309 |
| Female ID | Co-resident female receptiveness^(2)^ | Small swelling^(3)^ | 0.265 |
| Female ID | Co-resident female receptiveness^(2)^ | Medium swelling^(3)^ | -0.629 |
| Female ID | Co-resident female receptiveness^(2)^ | Male status^(4)^ | -0.264 |
| Female ID | Co-resident female receptiveness^(2)^ | Unit size^(1)^:Co-resident female receptiveness^(2)^ | -0.908 |
| Female ID | Co-resident female receptiveness^(2)^ | Unit size^(1)^:Small swelling^(3)^ | -0.776 |
| Female ID | Co-resident female receptiveness^(2)^ | Unit size^(1)^:Medium swelling ^(3)^ | -0.298 |
| Female ID | Small swelling^(3)^ | Medium swelling^(3)^ | 0.198 |
| Female ID | Small swelling^(3)^ | Male status^(4)^ | 0.247 |
| Female ID | Small swelling^(3)^ | Unit size^(1)^:Co-resident female receptiveness^(2)^ | -0.118 |
| Female ID | Small swelling^(3)^ | Unit size^(1)^:Small swelling ^(3)^ | -0.421 |
| Female ID | Small swelling^(3)^ | Unit size^(1)^:Medium swelling ^(3)^ | 0.470 |
| Female ID | Medium swelling^(3)^ | Male status^(4)^ | -0.278 |
| Female ID | Medium swelling^(3)^ | Unit size^(1)^:Co-resident female receptiveness^(2)^ | 0.728 |
| Female ID | Medium swelling^(3)^ | Unit size^(1)^:Small swelling^(3)^ | 0.790 |
| Female ID | Medium swelling^(3)^ | Unit size^(1)^:Medium swelling^(3)^ | -0.107 |
| Female ID | Male status^(4)^ | Unit size^(1)^:Co-resident female receptiveness^(2)^ | 0.023 |
| Female ID | Male status^(4)^ | Unit size^(1)^:Small swelling^(3)^ | -0.291 |
| Female ID | Male status^(4)^ | Unit size^(1)^:Medium swelling^(3)^ | 0.851 |
| Female ID | Unit size^(1)^:Co-resident female receptiveness^(2)^ | Unit size^(1)^:Small swelling^(3)^ | 0.711 |
| Female ID | Unit size^(1)^:Co-resident female receptiveness^(2)^ | Unit size^(1)^:Medium swelling^(3)^ | 0.274 |
| Female ID | Unit size^(1)^:Small swelling^(3)^ | Unit size^(1)^:Medium swelling^(3)^ | -0.345 |
| Male ID | Intercept | - | 0.631 |
| Male ID | Unit size^(1)^ | - | 0.394 |
| Male ID | Co-resident female receptiveness^(2)^ | - | 0.401 |
| Male ID | Small swelling^(3)^ | - | 0.518 |
| Male ID | Medium swelling^(3)^ | - | 0.205 |
| Male ID | Young adult female^(5)^ | - | 0.351 |
| Male ID | Mature adult female^(5)^ | - | 0.564 |
| Male ID | Old adult female^(5)^ | - | 1.010 |
| Male ID | Unit size^(1)^:Co-resident female receptiveness^(2)^ | - | 0.366 |
| Male ID | Unit size^(1)^:Small swelling^(3)^ | - | 0.174 |
| Male ID | Unit size^(1)^:Medium swelling^(3)^ | - | 0.238 |
| Male ID | Intercept | Unit size^(1)^ | -0.143 |
| Male ID | Intercept | Co-resident female receptiveness^(2)^ | -0.009 |
| Male ID | Intercept | Small swelling^(3)^ | -0.172 |
| Male ID | Intercept | Medium swelling^(3)^ | -0.359 |
| Male ID | Intercept | Young adult female^(5)^ | 0.888 |
| Male ID | Intercept | Mature adult female^(5)^ | 0.703 |
| Male ID | Intercept | Old adult female^(5)^ | 0.294 |
| Male ID | Intercept | Unit size^(1)^:Co-resident female receptiveness^(2)^ | -0.624 |
| Male ID | Intercept | Unit size^(1)^:Small swelling^(3)^ | -0.683 |
| Male ID | Intercept | Unit size^(1)^:Medium swelling^(3)^ | -0.524 |
| Male ID | Unit size^(1)^ | Co-resident female receptiveness^(2)^ | -0.897 |
| Male ID | Unit size^(1)^ | Small swelling^(3)^ | -0.614 |
| Male ID | Unit size^(1)^ | Medium swelling^(3)^ | -0.596 |
| Male ID | Unit size^(1)^ | Young adult female^(5)^ | 0.250 |
| Male ID | Unit size^(1)^ | Mature adult female^(5)^ | -0.191 |
| Male ID | Unit size^(1)^ | Old adult female^(5)^ | -0.088 |
| Male ID | Unit size^(1)^ | Unit size^(1)^:Co-resident female receptiveness^(2)^ | -0.557 |
| Male ID | Unit size^(1)^ | Unit size^(1)^:Small swelling^(3)^ | -0.623 |
| Male ID | Unit size^(1)^ | Unit size^(1)^:Medium swelling^(3)^ | -0.712 |
| Male ID | Co-resident female receptiveness^(2)^ | Small swelling^(3)^ | 0.894 |
| Male ID | Co-resident female receptiveness^(2)^ | Medium swelling^(3)^ | 0.852 |
| Male ID | Co-resident female receptiveness^(2)^ | Young adult female^(5)^ | -0.266 |
| Male ID | Co-resident female receptiveness^(2)^ | Mature adult female^(5)^ | -0.125 |
| Male ID | Co-resident female receptiveness^(2)^ | Old adult female^(5)^ | -0.279 |
| Male ID | Co-resident female receptiveness^(2)^ | Unit size^(1)^:Co-resident female receptiveness^(2)^ | 0.419 |
| Male ID | Co-resident female receptiveness^(2)^ | Unit size^(1)^:Small swelling^(3)^ | 0.674 |
| Male ID | Co-resident female receptiveness^(2)^ | Unit size^(1)^:Medium swelling^(3)^ | 0.742 |
| Male ID | Small swelling^(3)^ | Medium swelling^(3)^ | 0.967 |
| Male ID | Small swelling^(3)^ | Young adult female^(5)^ | -0.267 |
| Male ID | Small swelling^(3)^ | Mature adult female^(5)^ | -0.484 |
| Male ID | Small swelling^(3)^ | Old adult female^(5)^ | -0.655 |
| Male ID | Small swelling^(3)^ | Unit size^(1)^:Co-resident female receptiveness^(2)^ | 0.218 |
| Male ID | Small swelling^(3)^ | Unit size^(1)^:Small swelling^(3)^ | 0.596 |
| Male ID | Small swelling^(3)^ | Unit size^(1)^:Medium swelling^(3)^ | 0.589 |
| Male ID | Medium swelling^(3)^ | Young adult female^(5)^ | -0.481 |
| Male ID | Medium swelling^(3)^ | Mature adult female^(5)^ | -0.623 |
| Male ID | Medium swelling^(3)^ | Old adult female^(5)^ | -0.692 |
| Male ID | Medium swelling^(3)^ | Unit size^(1)^:Co-resident female receptiveness^(2)^ | 0.393 |
| Male ID | Medium swelling^(3)^ | Unit size^(1)^:Small swelling^(3)^ | 0.724 |
| Male ID | Medium swelling^(3)^ | Unit size^(1)^:Medium swelling^(3)^ | 0.650 |
| Male ID | Young adult female^(5)^ | Mature adult female^(5)^ | 0.567 |
| Male ID | Young adult female^(5)^ | Old adult female^(5)^ | 0.173 |
| Male ID | Young adult female^(5)^ | Unit size^(1)^:Co-resident female receptiveness^(2)^ | -0.895 |
| Male ID | Young adult female^(5)^ | Unit size^(1)^:Small swelling^(3)^ | -0.877 |
| Male ID | Young adult female^(5)^ | Unit size^(1)^:Medium swelling^(3)^ | -0.715 |
| Male ID | Mature adult female^(5)^ | Old adult female^(5)^ | 0.875 |
| Male ID | Mature adult female^(5)^ | Unit size^(1)^:Co-resident female receptiveness^(2)^ | -0.181 |
| Male ID | Mature adult female^(5)^ | Unit size^(1)^:Small swelling^(3)^ | -0.406 |
| Male ID | Mature adult female^(5)^ | Unit size^(1)^:Medium swelling^(3)^ | -0.157 |
| Male ID | Old adult female^(5)^ | Unit size^(1)^:Co-resident female receptiveness^(2)^ | 0.163 |
| Male ID | Old adult female^(5)^ | Unit size^(1)^:Small swelling^(3)^ | -0.163 |
| Male ID | Old adult female^(5)^ | Unit size^(1)^:Medium swelling^(3)^ | 0.030 |
| Male ID | Unit size^(1)^:Co-resident female receptiveness^(2)^ | Unit size^(1)^:Small swelling^(3)^ | 0.896 |
| Male ID | Unit size^(1)^:Co-resident female receptiveness^(2)^ | Unit size^(1)^:Medium swelling^(3)^ | 0.827 |
| Male ID | Unit size^(1)^:Small swelling^(3)^ | Unit size^(1)^:Medium swelling^(3)^ | 0.947 |

^(1)^ z-transformed to a mean of zero and a standard deviation (sd) of one; mean and sd of the original variable were 3.14 and 1.72, respectively

^(2)^ centred and dummy coded with no being the reference level

^(3)^ centred and dummy coded with large swelling being the reference level

^(4)^ centred and dummy coded with primary male being the reference level

^(5)^ centred and dummy coded with subadult female being the reference level

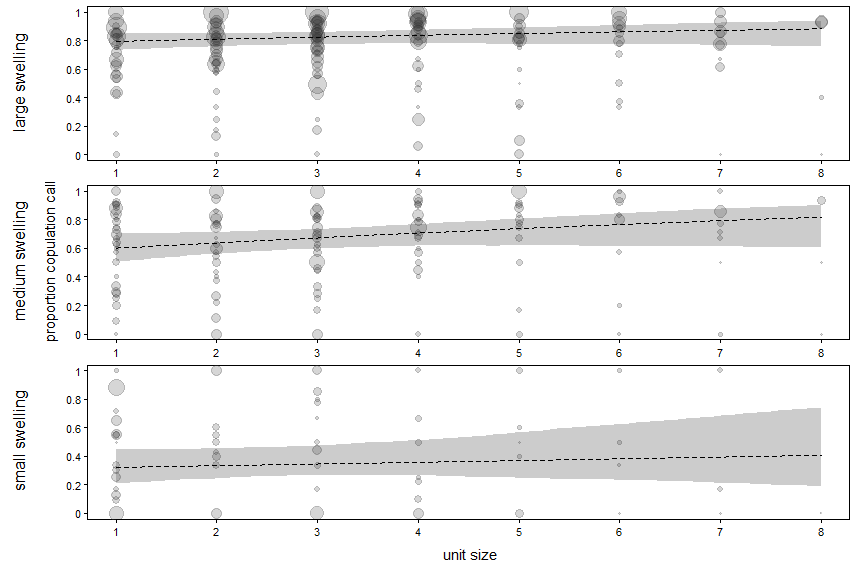

**Figure S1.** Proportion of copulation calls given in relation to unit size and swelling size. Indicated are the fitted model (dashed lines) and its 95% confidence intervals (grey polygons) with all other terms in the model being at their average. Dots represent the proportion of copulations with calls for each combination of unit size and female ID. The area of the dots is proportional to the number of copulations per female and unit size (range: 1 – 72).

**Table S2.** Results of the reduced model of the effect of unit size on the probability to produce copulation calls (estimates together with standard errors, confidence limits, and significance test)

| **Term** | **Estimate** | **SE** | **Lower CI** | **Upper CI** | **X2** | **df** | **P** |
| --- | --- | --- | --- | --- | --- | --- | --- |
| Intercept | 1.554 | 0.186 | 1.210 | 1.910 |  |  |  |
| Unit size^(1)^ | 0.192 | 0.116 | -0.049 | 0.441 | 2.629 | 1 | 0.105 |
| Co-resident female receptiveness^(2)^ | -0.114 | 0.168 | -0.400 | 0.201 | 0.453 | 1 | 0.501 |
| Small swelling^(3),(4)^ | -2.213 | 0.200 | -2.659 | -1.846 | 64.992 | 2 | <0.001 |
| Medium swelling^(3)^ | -0.804 | 0.122 | -1.020 | -0.572 |  |  |  |
| Young adult female^(5),(6)^ | -0.254 | 0.217 | -0.669 | 0.197 | 4.726 | 3 | 0.193 |
| Mature adult female^(5)^ | -0.014 | 0.213 | -0.442 | 0.386 |  |  |  |
| Old adult female^(5)^ | 0.612 | 0.376 | -0.041 | 1.354 |  |  |  |
| Male status^(7)^ | -0.222 | 0.196 | -0.602 | 0.202 | 1.279 | 1 | 0.258 |

^(1)^ z-transformed to a mean of zero and a standard deviation (sd) of one; mean and sd of the original variable were 3.14 and 1.72, respectively

^(2)^ dummy coded with no being the reference level

^(3)^ dummy coded with large swelling being the reference level

^(4)^ the indicated test refers to the overall effect of swelling size

^(5)^ dummy coded with subadult female being the reference level

^(6)^ the indicated test refers to the overall effect of female age category

^(7)^ dummy coded with primary male being the reference level
